# *netSmooth*: Network-smoothing based imputation for single cell RNA-seq

**DOI:** 10.1101/234021

**Authors:** Jonathan Ronen, Altuna Akalin

## Abstract

Single cell RNA-seq (scRNA-seq) experiments suffer from a range of characteristic technical biases, such as dropouts (zero or near zero counts) and high variance. Current analysis methods rely on imputing missing values by various means of local averaging or regression, often amplifying biases inherent in the data. We present netSmooth, a network-diffusion based method that uses priors for the covariance structure of gene expression profiles on scRNA-seq experiments in order to smooth expression values. We demonstrate that netSmooth improves clustering results of scRNA-seq experiments from distinct cell populations, time-course experiments, and cancer genomics. We provide an R package for our method, available at: https://github.com/BIMSBbioinfo/netSmooth.

## Introduction

Single cell RNA sequencing (scRNA-seq) enables profiling of single cells’ transcriptomes at unprecedented throughput and resolution. It has enabled previously impractical, studies of cell type heterogeneity, differentiation, and developmental trajectories [1]. However, the adaptation of RNA sequencing techniques from bulk samples to single cells did not progress without challenges. Typically, only a fraction of a cells transcriptome may be captured by the experiment, leading to so called “drop-out” events where a gene gets a false 0 (or near 0) count in some cell. The dropout rate is related to the population level expression of a gene leading to many false zero counts for lowly expressed genes, and artificially low counts for highly expressed ones [2]. Furthermore, the drop-out rate could be related to the biology of the cell type, as some cell types transcribe fewer genes than others, which will appear as drop-out events [2]. When summed over many samples, transcript counts from single cells resemble those of bulk experiments [3], but across individual cells there is significant variation. This makes analysis more difficult than in bulk RNA sequencing experiments.

Computational methods designed to deal with these issues treat dropout events as missing data points, whose values may be imputed based on non-missing data points (observed measurements). The proportion of 0 counts per gene, a proxy for its technical dropout rate, is a function of the population-wise mean expression of that gene [4, 2]. This observation has led researchers to treat 0 counts as dropout candidates to be imputed.

CIDR [5] attempts to impute missing values based on the predicted mean expression of a gene, given its empirical dropout rate (0-count). scImpute [6] estimates dropout likelihoods per gene and per sample, and assigns each gene in each sample a status as a dropout candidate. Genes might be considered likely dropouts even with nonzero expression, and 0-count genes might not be considered likely dropouts, based on their population-wide expression distributions. It then uses a regularized linear model to predict the expression of dropout genes based on the expression of likely non-dropouts in all other cells. MAGIC [7] performs local averaging after building a topological graph of the data, updating the expression value of all genes in all cells to their local neighborhood average.

All of the methods mentioned above use measured information in the data in order to impute the missing information within the same data. As such, they amplify whatever biases are present in a dataset; similar cells preimputation will become more similar after imputation, as expression profiles of non-dropout genes will drive similarities in imputed dropped-out genes. Further, all methods except MAGIC only impute unobserved expression events (0s or near 0s), while the dropout phenomenon actually affects the whole transcriptome. Hence, imputation methods for scRNAseq should also adjust non-0 expression measurements in order to recover the true signal. We present a method, called *netSmooth*, that uses prior knowledge to temper noisy experimental data. RNA sequencing experiments produce counts data as a proxy for gene activity, which is not known a-priori, especially for experiments profiling unknown cell types. However, decades of molecular biology research have taught us much about the principles of gene interaction. Interacting genes are likely to be co-expressed in cells [8, 9], and as such, protein-protein interaction (PPI) databases [10, 11] describe genes’ propensity for co-expression. We developed a graph-diffusion method on PPI networks for smoothing of gene expression values. Each node in the graph (a gene) has an associated gene expression value, and the diffusion presents a weighted averaging of gene expression values among adjacent nodes in the graph, within each cell. This is done iteratively until convergence, strengthening co-expression patterns which are expected to be present. Incorporation of prior data from countless experiments in the preprocessing of scRNA-seq experiments improves resistance to noise and dropouts. Similar network based approaches have been used to extract meaningful information from sparse mutational profiles [12, 13], and indirectly on gene expression data by diffusing test statistics on the network to discover regulated gene candidates [14]. We propose diffusion of gene expression values directly on the network as a method for data denoising and imputation. Furthermore, the parameters of this proposed method could be optimized using clustering robustness metrics. We applied our method to a variety of single cell experiments and compared its performance to other selected imputation methods scImpute and MAGIC. These methods represent the latest and divergent ways of imputing the scRNA-seq data.

We also made available an R package providing the necessary functionality to use our method on other data. It is available on GitHub: https://github.com/BIMSBbioinfo/netSmooth.

## Results

### Overview of the method

The intuition behind the *netSmooth* algorithm is that gene networks encoding co-expression patterns can be used to smooth scRNA-seq data, pushing its coexpression patterns in a biologically meaningful direction. We demonstrate this using protein-protein interaction networks, which are predictive of coexpression [9]. We produced a PPI graph of high-confidence interactions based on the PPI database STRING [10].

There are 2 inputs to the method: (1) a gene expression matrix, *N* genes by *M* cells, and (2) a graph where genes are nodes, and edges indicate genes which are expected to be co-expressed. The edges may be weighed, indicating the strength or direction of a relationship; an edge weight of 2 indicates stronger expected co-expression than an edge weight of 1, and an edge weight of–1 indicates negative expected co-expression, such as one gene being a repressor for another. The expression profile of each cell is then projected onto the graph, and a diffusion process is used to smooth the expression values, within each sample, of adjacent genes in the graph (Figure 1). In this way, post-smoothing values of genes represent an estimate of activity levels based on reads aligned to that gene, as well as those aligned to its neighbors in the graph. Thus, a gene with a low read count (possible technical drop-out), whose neighbors in the graph are highly expressed, will get a higher value post smoothing. The rate at which expression values of genes diffuse to their neighbors is degree-normalized, so that genes with many edges will affect their neighbors less than genes with more specific interactions. The diffusion is done using a “random walks with restarts” (RWR) process [13], where a conceptual random walker starts in some node in the graph, and at each iteration moves to a neighboring node with a probability determined by the edge weight between the nodes, or, with some probability, restarts the walk from the original node. The *network-smoothed* value is the stationary distribution of this process. The RWR process has one free parameter, the restart rate. A low value for the restart rate allows diffusion to reach further in the graph; a high restart rate will lead to more local diffusions. For more details see the Methods section.

**Figure 1.**
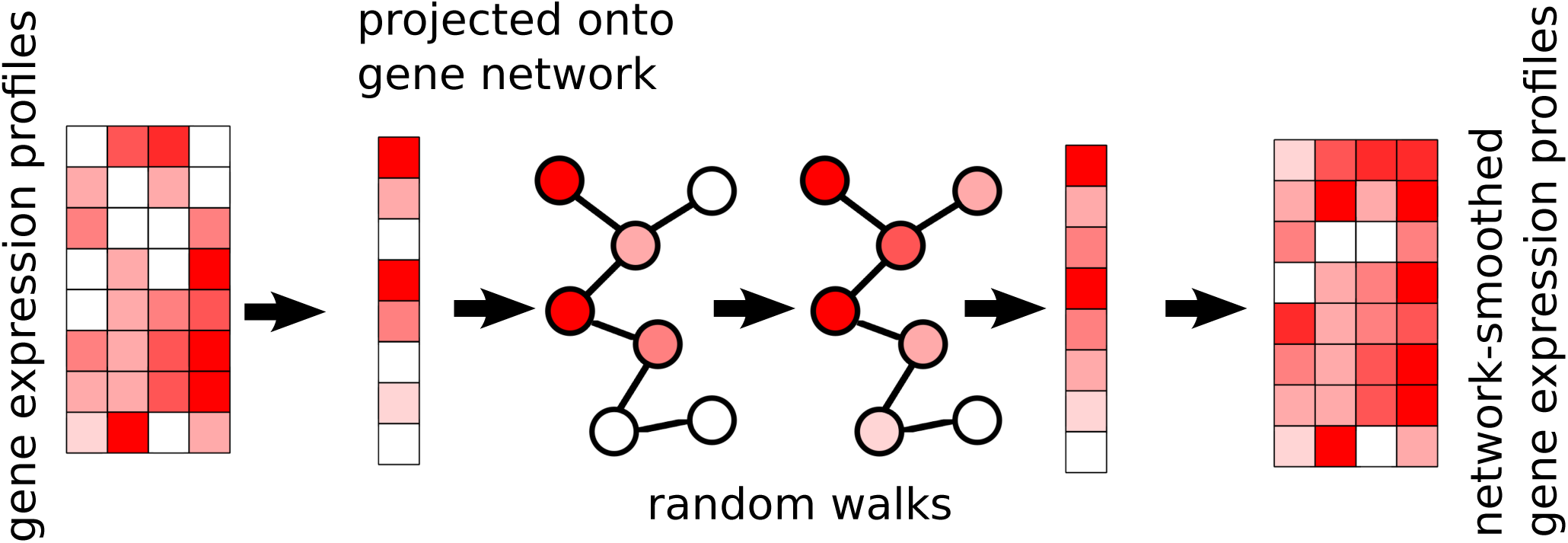
The *netSmooth* algorithm takes a gene expression profile, and a gene network. The expression profile of each sample is projected onto the network, where a diffusion process allows genes’ expression values to be smoothed by their neighbors’. This is done for each cell independently of others. The end result is a network smoothed gene expression matrix.

### Network smoothing improves cell type identification from single-cell RNA-seq

We first assess *netSmooth* on a dataset of 1645 mouse hematopoietic stem/progenitor cells (HSPCs) assayed using flow cytometry as well as scRNA-seq [15]. The cells are FACS-sorted into 12 common HSPC phenotypes. This presents an atlas of the hematopoiesis process at a single cell resolution, showing the differentiation paths taken by E-SLAM HSCs as they differentiate to E, GM, and L progenitors. The authors of this study demonstrate that upon clustering the data, some clusters corresponds to cell types. However, the clusters are not noise free and do not fully recapitulate cell type identity. We obtained clusterings of the cells from the normalized counts, as well as after application of *netSmooth*, MAGIC [7], and scImpute [6], using a robust clustering procedure based on the *clusterExperiment* R package [16] (See Methods). After clustering, we used the edgeR-QLF test [17] to identify genes that are differentially expressed in any of the discovered clusters. Figure 2a,b shows that after network-smoothing, we are able to identify clusters with a more pronounced differential expression profile. Further, many more of the genes identified as differentially expressed between the clusters (without smoothing) seem to have low and uninformative expression values overall. MAGIC and scImpute also improve this pattern (Figure 2c,d). MAGIC seems to do the strongest transformation to the data, as seen in lower dimension embeddings (Figures S2, S3).

**Figure 2.**
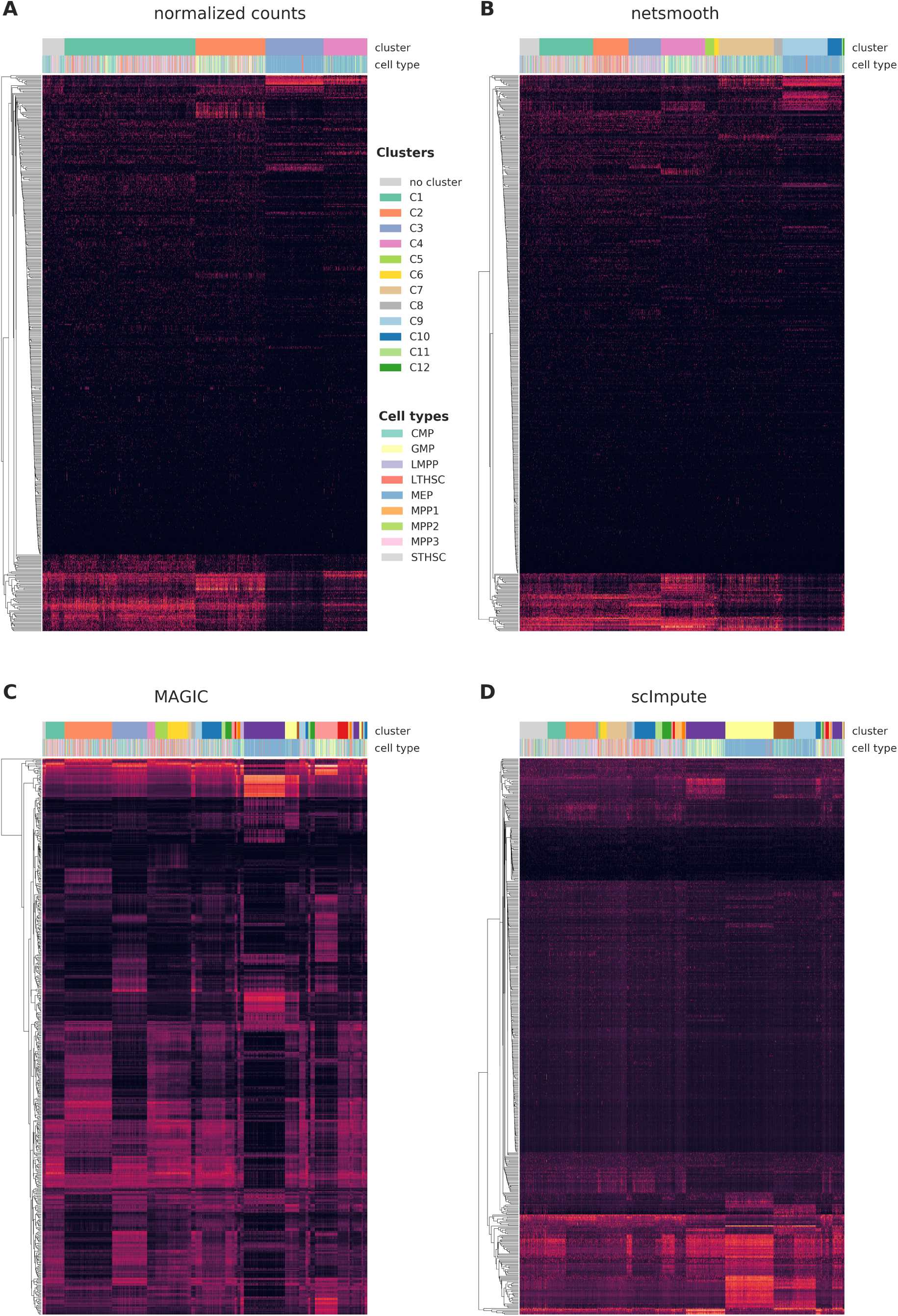
Cells were clustered using the robust clustering procedure, and the 500 most differentially expressed genes (by edgeR-QLF test adjusted P value) in any of the discovered clusters are shown in a heatmap, as well as cluster assignments and FACS-sorted cell types. A) raw (no imputation), B) after application of *netSmooth*, C) missing values imputed using MAGIC D) missing values imputed using scImpute.

As this dataset has cells with labels independent of the RNAseq (FACS-sorted phenotypes), it presents us with an opportunity to compare the gene expression levels (as measured by RNAseq), to a meaningful phenotypic variable, i.e. the cell type. The cell type discrimination of a clustering result is compared using a cluster purity metric and and the adjusted mutual information (AMI). The cluster purity measures how cell-type specific clusters are by comparing homogeneity of the external labels (FACS-defined cell types), within clusters provided by scRNA-seq data. AMI is a chance-adjusted information-theoretic measure of agreement between two labellings. This method accounts for artificially high mutual information between external labels and clusters when there are high number of clusters (See Methods for details on metrics). We also measured number of cells in robust clusters as quantitative metric. The robust clustering procedure allows cells to be omitted (not be assigned to a cluster) if they cannot be placed in a cluster across multiple clustering methods and/or parameters (See Methods). Only MAGIC is able to increase the proportion of cells in this dataset which fall into robust clusters (Figure 3a), but only *netSmooth* leads to more biologically meaningful clusters, in terms of purity and AMI (Figures 3b,c), demonstrating that *netSmooth* can assist in cell type identification, and outperformed both MAGIC and scImpute in this task. The higher clusterability following application of MAGIC than *netSmooth*, might indicate that MAGIC was overzealous in its transformation, squeezing more cells into the same space. This might lead to more robust clusters, but less reliable cell type identification.

**Figure 3.**
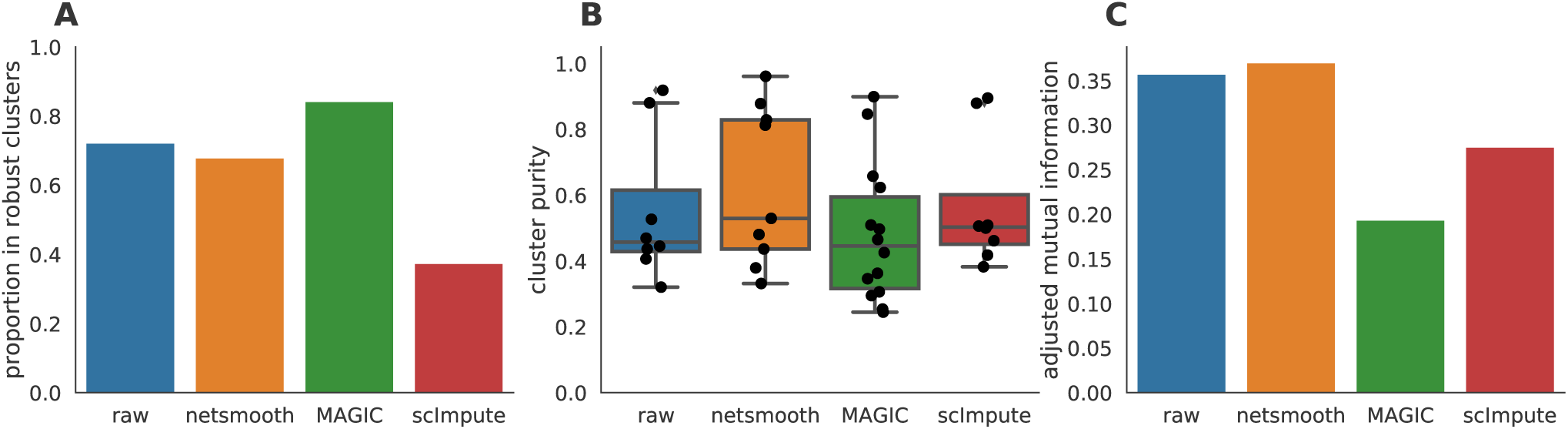
A) The proportion of cells which were assigned to robust clusters. B) cluster purity (proportion of dominant cell type) for the robust clusters. *netSmooth* produces the most pure clusters in terms of cell types. C) AMI of the clustering results obtained after application of each of the methods. Only *netSmooth* increases the AMI between the clustering and the cell types.

### Network smoothing improves capture of developmental expression patterns

Next, we test *netSmooth* on 269 isolated cells from mouse embryos at different stages of pre-implantation development between oocyte and blastocyst, as well as 5 liver cells and 10 fibroblast cells [18]. The authors of this study demonstrated that lower dimension embeddings capture much of the developmental trajectory (Figure 4a, S5a, Figure S4a). We then applied *netSmooth*, MAGIC, and scImpute. Figure 4b shows the principal component analysis of *netSmooth*-processed data, and Figures 4c and 4d show the PCA plot following application of MAGIC and scImpute, respectively. *netSmooth* and scImpute preserve most of the variance structure of the data, while MAGIC seems to push the data onto a completely different manifold (Figure 4, Figure S5). We used the robust clustering procedure to obtain clusters, and computed the cluster purity and AMI metrics. *netSmooth* enabled the clustering procedure to place more of the samples into robust clusters (Figure 5a), and as in the hematopoiesis case, *netSmooth* is able to assist in identifying the developmental stage or tissue that cells belong to better than the other methods, as evidenced by the higher cluster purities (Figure 5b) and AMI (Figure 5c). Although MAGIC and scImpute reduce the 0-count genes further than *netSmooth* (Figure S1), they do not add as much clarity to the developmental stage signal inherent in the data. This shows that imputing missing counts based on data from the same experiment is not as powerful as including priors in the quasi-imputation process *netSmooth* does.

**Figure 4.**
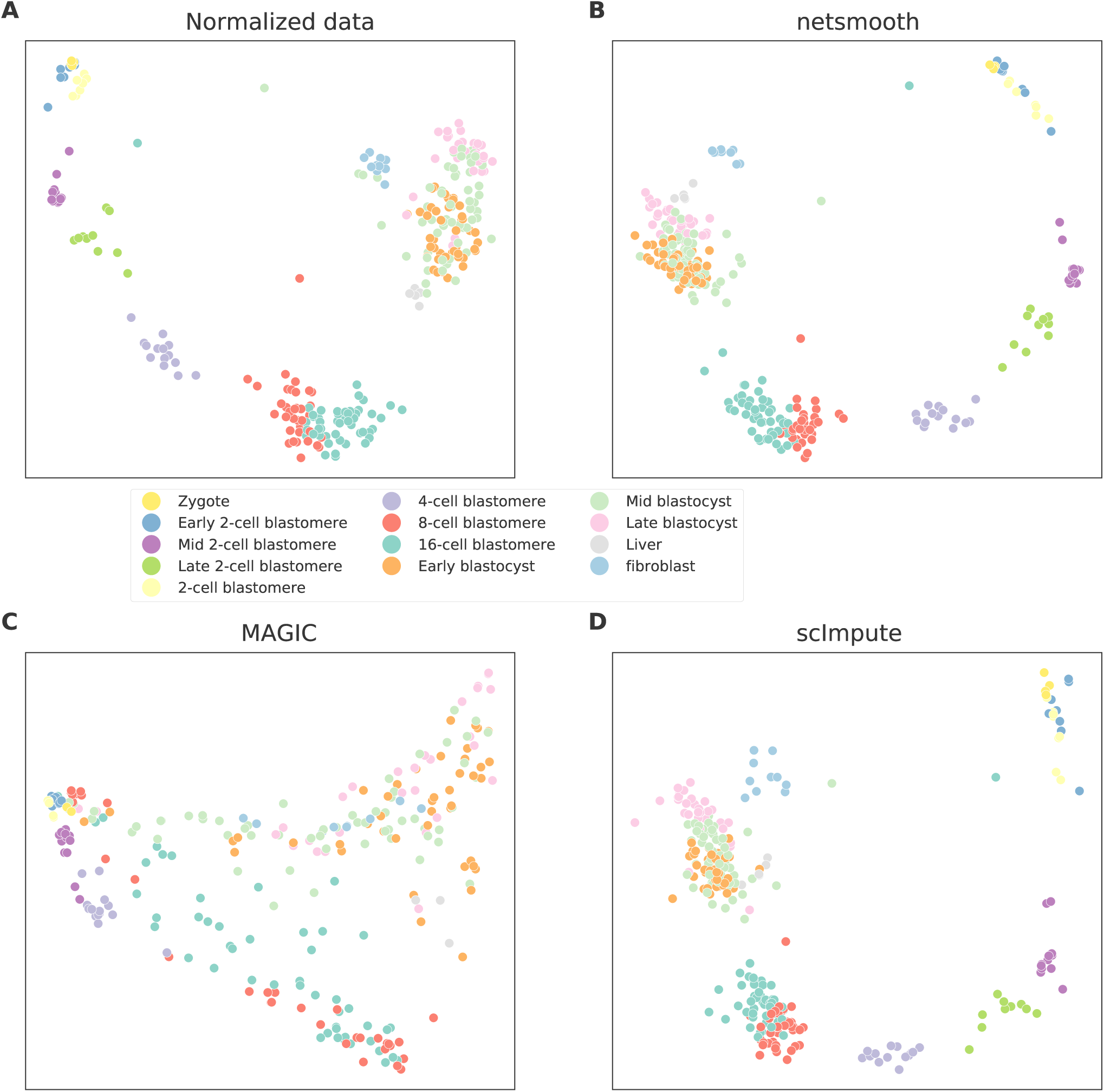
2D PCA plots of the embryonic development dataset A) no preprocessing, B) after application of *netSmooth*, C) after imputing missing values with scImpute, and D) after application of MAGIC.

**Figure 5.**
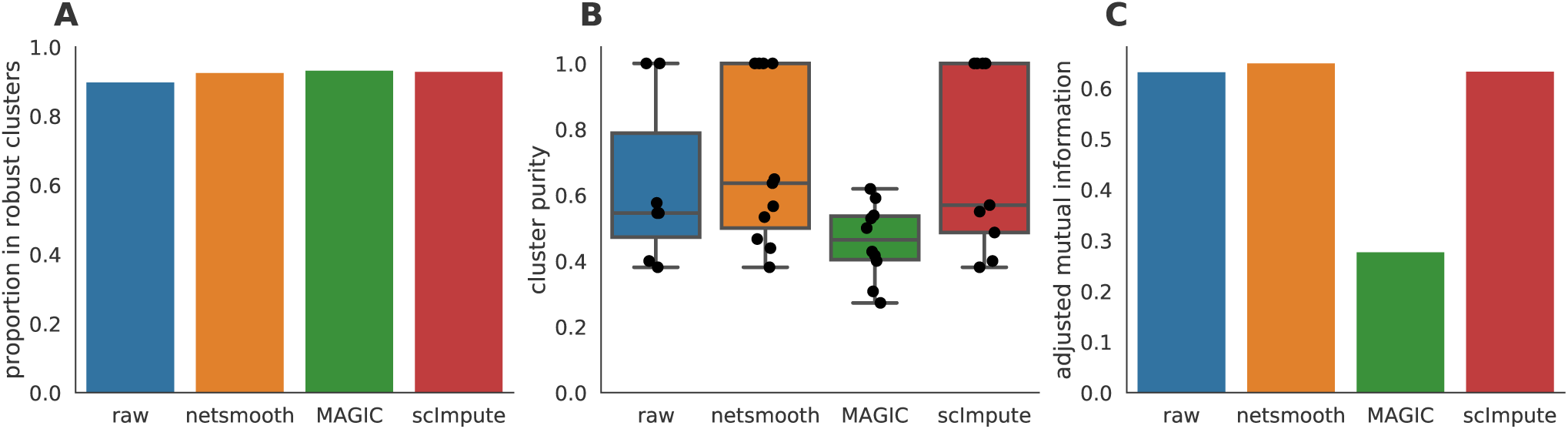
The Embryonic development dataset. A) The proportion of cells which were assigned to robust clusters. All three methods lead to better clusterability, with MAGIC having the strongest effect. B) cluster purity (proportion of dominant cell type) for the robust clusters. *netSmooth* produces the most pure clusters in terms of cell types. C) Adjusted mutual information of clusterings and cell types. Only *netSmooth* increases the AMI over the non-preprocessed data.

### Network smoothing improves identification of glioblastoma tumors

Finally, we demonstrate applicability of *netSmooth* to cancer research. Patel et al. generated scRNA-seq data of 800 cells from 5 glioblastoma tumors and 2 cell lines [19]. Lower dimension embedding plots show that cells from different tumors or cell lines generally group together, but some are not wholly distinguishable from other tumors (Figure 6a, S7a, S6a). Further, the two cell lines group closer to each other than the other patient samples. After applying *netSmooth* to the data, tumors become easier to distinguish in a lower dimensional embedding (Figure 6b), indicating that *netSmooth* improves assignment of each cell to its tumor, cell line, or clone of origin. Again, scImpute also leads to similar reduced dimension embedding (Figure 6d), while MAGIC distorted the data more than the other methods (Figure 6c). We used the robust clustering procedure before and after *netSmooth*, MAGIC, and scImpute. Only MAGIC increase the clusterabitliy of the data (Figure 7a), but *netSmooth* leads to the most pure clusters, in terms of tumor or cell line of origin (Figure 7b, Figure 7c).

**Figure 6.**
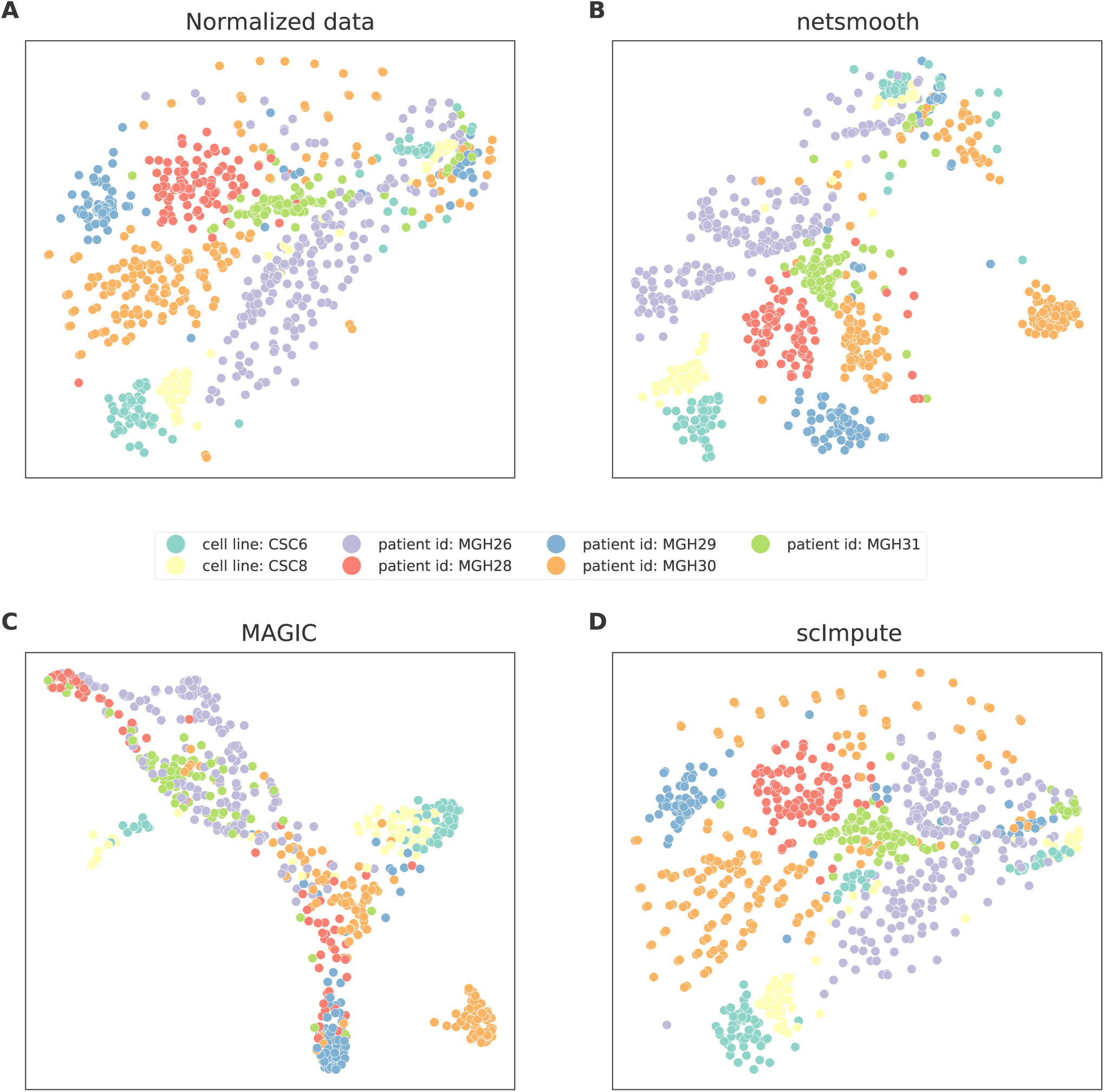
t-SNE plots of the glioblastoma dataset A) no preprocessing, B) after application of *netSmooth*, C), using MAGIC, and D) after application of scImpute.

**Figure 7.**
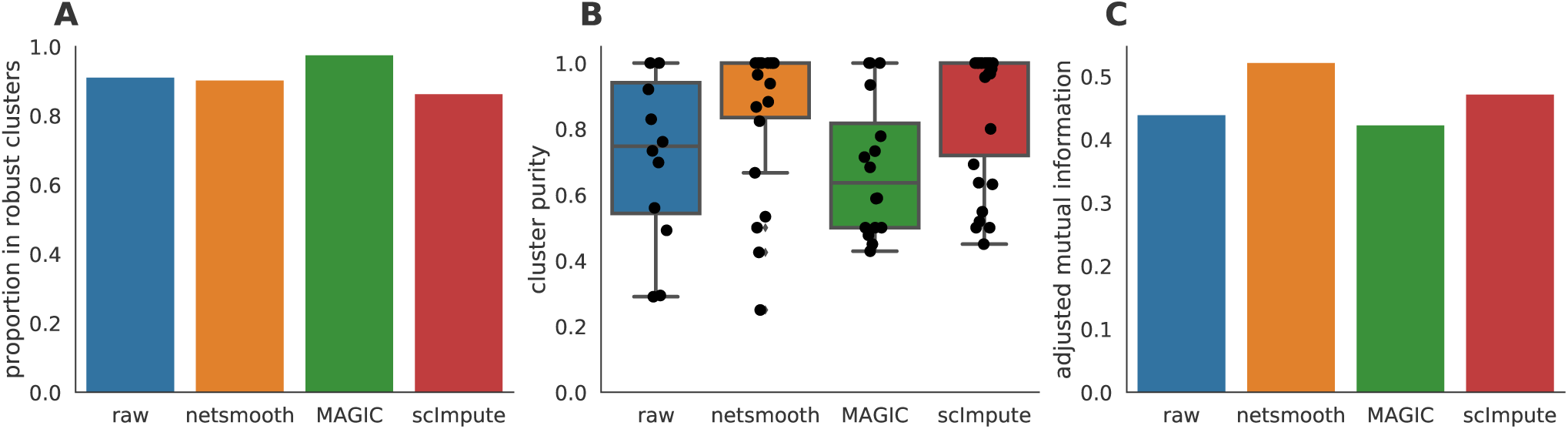
Imputation performance for the glioblastoma dataset. A) The proportion of cells which were assigned to robust clusters. *netSmooth*, MAGIC, and scImpute all increased the proportion of cells that are assigned to robust clusters, with MAGIC leading, *netSmooth* in second place, and scImpute in third. B) cluster purity (proportion of dominant cell type) for the robust clusters. *netSmooth* produces the most pure clusters in terms of tumor or cell line of origin. C) AMI of the clustering results obtained after application of each of the methods.

Tumor or cell line of origin is an imperfect proxy for phenotypical variation in cancer cells, because some cells cluster by cell type rather than tumor of origin, demonstrating the heterogeneity in these glioblastoma tumors and similarities across origins [19]. Nevertheless, we chose to compute cluster purity based on the cell origin rather than other labels which might be assigned to them, as it is the only *ground truth* variable that is independent of the RNAseq experiment. Further, cells do group by origin (Figure 6, Figure S6), and identification of origin is an interesting question in its own right in the field of cancer genomics, particularly for heterogeneous tumors such as these.

### Sensitivity to the network

Next, we set out to ensure that the results are not an artifact of the network structure, i.e. that the actual links between genes that we used in the network are important. We expect *netSmooth* not to perform well when using networks with similar characteristics, but where edges do not represent real interactions. To that effect, we constructed 20 random networks by keeping the same graph structure of the real PPI graph, but shuffling the gene names.

Thus, these random networks share all the characteristics of the real network (degree distribution, community structure), except for the true identity of the nodes. We then used those networks as inputs to *netSmooth* and ran the benchmarks as before on the hematopoiesis dataset. Using random networks as an input to *netSmooth* gives cluster purities distributed around a mode given by the cluster purities of the raw data, while the cluster purities given from using the real PPI network lie at the extreme edge of the distribution (Figure 8a). Further, most random networks result in fewer samples belonging to robust clusters (Figure 8b). These results demonstrate that it is indeed the information contained in the PPI graph enables netSmooth to transform the gene expression matrix in a more biologically coherent direction, and that the transformation we see can not be explained simply by the network structure.

**Figure 8.**
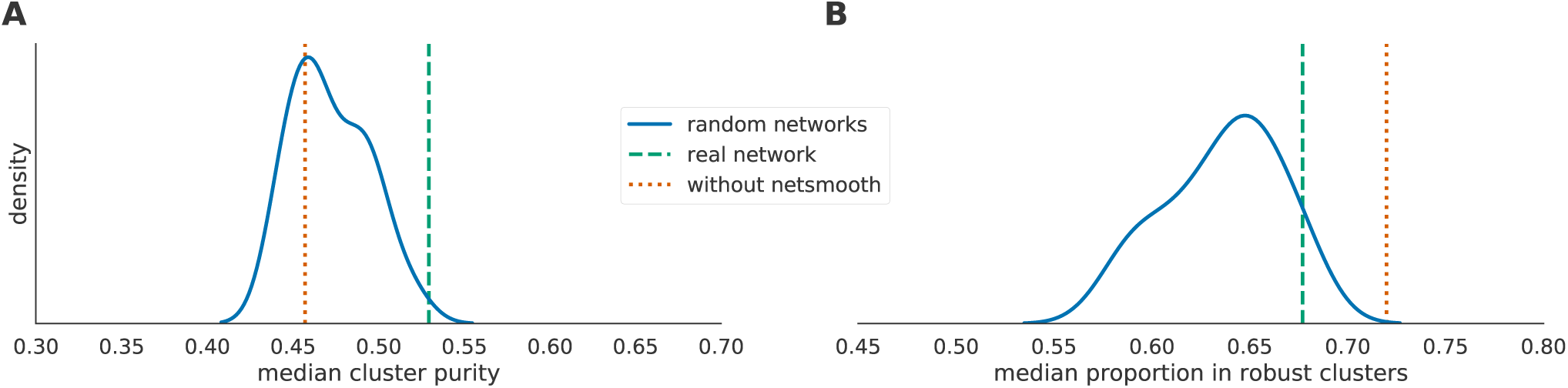
Performance of *netSmooth* with randomized networks. A) The median cluster purity achieved with the random networks. The real network outperforms the random ones, which result in cluster purities distributed around the purity given without using *netSmooth*. B) The proportion of samples assigned to robust clusters using the random networks as well as the real one. While all networks result in fewer samples robustly clustered (in the hematopoiesis dataset), the real network outperforms most random networks.

### Using other networks with netSmooth

In addition to using an unweighed (where all edge weights are 1), undirected (where all edge weights are positive) network from string-db, we constructed other gene networks and used them as inputs to *netSmooth*. We created a directed gene network from only those edges in string-db which are marked as activating or inhibiting^1^. We set the edge weights of the activating interactions to +1, and –1 for the inhibiting interactions, allowing gene expression values to be adjusted downwards for genes whose known antagonists are highly expressed. After smoothing, we set all negative smoothed expression values to 0. We also constructed a gene network from string-db using only genes that are known to demonstrate cell-type specific expression. In order to obtain a list of genes with such cell-type specific expression patterns from the *Expression Atlas* [20], we used only the genes which show a cell-type specific expression with a mean TPM of at least 1 in some cell type, and used the subset of string-db network containing those genes as an input to *netSmooth.* Both of those modified graphs perform similarly to the undirected graph from string-db (Figure 9, Figure S8a, Figure S8b), demonstrating that *netSmooth* is able to use priors from different types of experiments in order to improve clustering of scRNA-seq.

**Figure 9.**
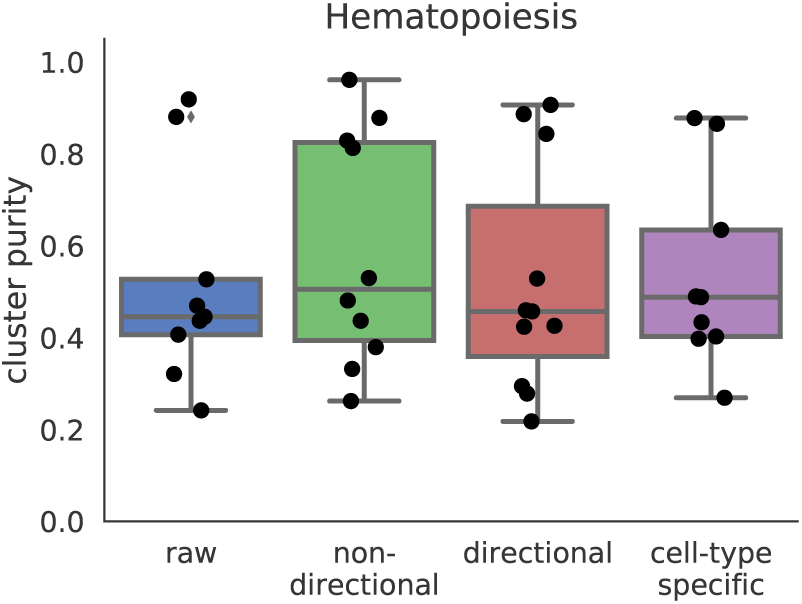
Cluster purities after applying *netSmooth* with different input networks. Raw refers to no smoothing, non-directional is the same as the results shown in previous sections. Directional refers to a gene network where inhibitory relationships have negative edge weights, and cell-type specific refers to a gene network of only genes which are known to have cell-type specific expression patterns.

We also considered other sources for the gene network. We constructed a gene network from HumanNet [21], a functional gene network where edges denote interactions between two genes. We constructed a smoothing graph by taking all edges from HumanNet, and producing a graph where all edge weights are set to 1. We then used this graph as an input to *netSmooth* on the glioblastoma dataset. It performs similarly to the network from string-db (Figure 10, Figure S8c), demonstrating that other sources for gene interactions may also be used by *netSmooth* to improve clustering results of scRNA-seq.

**Figure 10.**
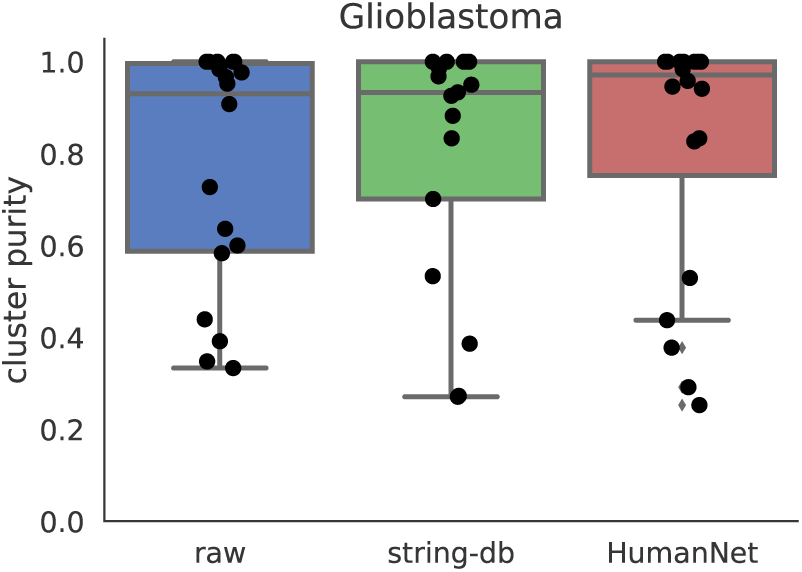
Cluster purities after applying *netSmooth* with different input networks. Raw refers to no smoothing, string-db is the same as the results shown in previous sections, and HumanNet refers to a gene network constructed from the HumanNet database.

### Optimizing the smoothing parameters by cluster robustness

The *netSmooth* algorithm, given a gene network, has one free parameter - the restart rate of the random walker, (1 – *α*). Alternatively, *α* is the complement of the restart rate. An *α* = 0 indicates a perfect restart rate and consequently no smoothing; an *α* = 1 corresponds to a random walk without restarts. Intermediate values for *α* result in increasing levels of smoothing; the value of *α* determines how far random walks will go on the graph before restarting, or how far along the network a gene’s influence is allowed to reach (See Methods). It is tempting to optimize *α* with respect to the variable the experiment sets out to measure, e.g. cluster purity. For instance, in the embryonic development dataset, we would choose *α* = 0.7 as the value that produces the highest cluster purity (Figure 11b). However, in many experiments the identity of the samples is not known a-priori. Therefore, we propose a data driven workflow to pick a sensible value for *α*.

**Figure 11.**
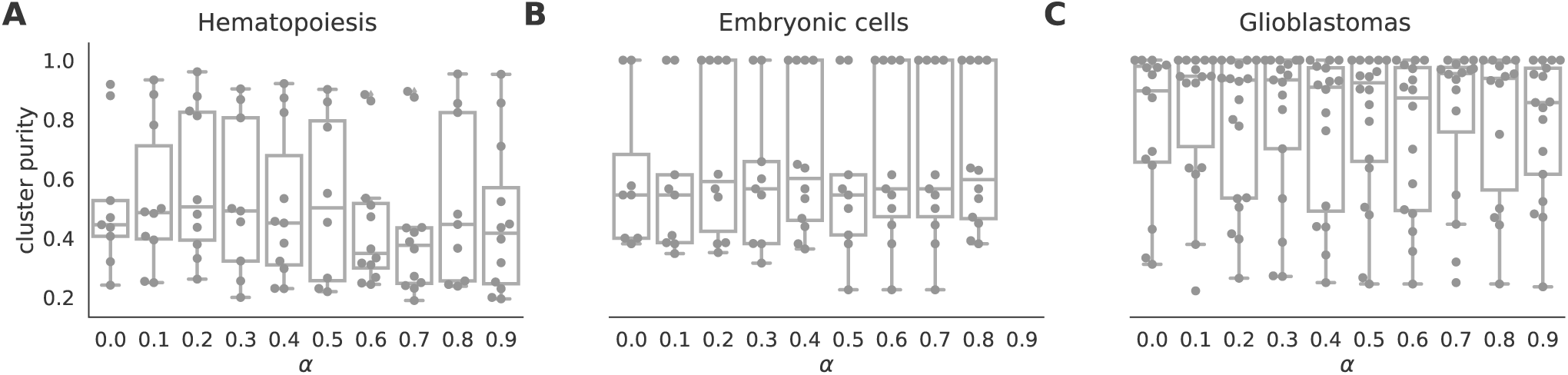
boxplots of cluster purity for clusters obtained by the robust clustering procedure following application of *netSmooth* with different values of *α*. *α* = 0 is equivalent to not using *netSmooth* at all. The procedure is robust to alpha, that is, most values of alpha produce more robust clusters. A) HSPCs, B) embryonic cells, C) glioblastomas.

One such data-driven statistic is the proportion of samples assigned to robust clusters; following application of *netSmooth*, the robust clustering procedure is able to assign more samples to statistically robust clusters. For all three datasets, picking the *α* that gives the highest proportion of cells in robust clusters, also gives the clusters with the highest purity index (Figure 12). Importantly, this metric is entirely data-driven and does not require external labels, making it feasible for any scRNA-seq study. The results in the previous sections all use the value of *α* picked to optimize proportion in robust clusters.

**Figure 12.**
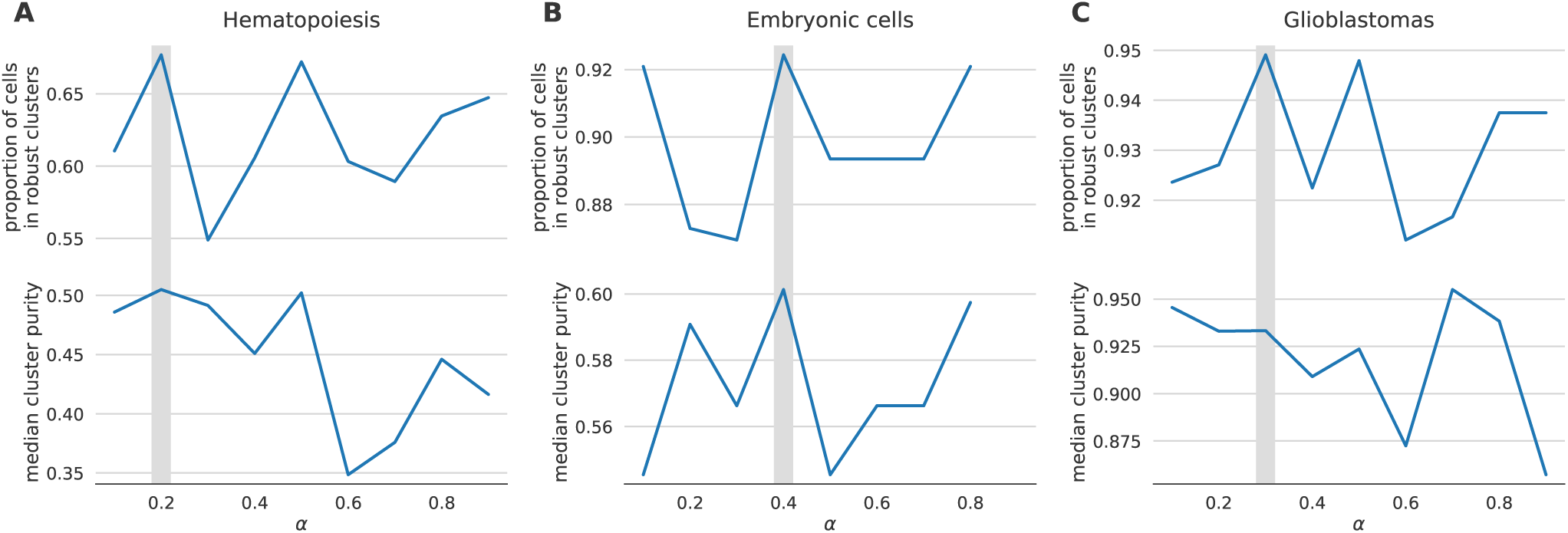
the proportion of cells in robust clusters, and cluster purity for those robust clusters, for a range of alpha values, shows that picking the alpha with the highest proportion in robust clusters also picks the alpha with the highest cluster purity. A) hematopoietic stem/progenitor cells B) embryonic cells, C) glioblastomas.

## Discussion

Single cell RNA sequencing technology provides whole-genome transcriptional profiles at unprecedented throughput and resolution. However, high variance and dropout events that happen in all current scRNA-seq platforms complicate the interpretation of the data. Methods that treat 0 counts as missing values and impute them based on nonzero values in the data may amplify biases in the data.

We presented *netSmooth* as a preprocessing step for scRNA-seq experiments, overcoming these challenges by the use of prior information derived from protein-protein interactions or other molecular interaction networks. We demonstrated that network smoothing assists in several standard analyses that are common in scRNA-seq studies. This procedure enhances cell type identification in hematopoiesis; it elucidates time series data and assists identification of the developmental stage of single cells. Finally, it is also applicable in cancer, improving identification of tumor of origin for glioblastomas. In addition, we showed that network smoothing parameter can be optimized by cluster robustness metrics, providing a workflow when there are no other external labels to distinguish cells. We demonstrated that *netSmooth* can use prior information from different sources in order to achieve this. We compared *netSmooth* with scImpute, a statistical genome-wide imputation method, and MAGIC, a genome-wide data smoothing algorithm, and demonstrated that while scImpute and MAGIC reduce the drop-out phenomenon more than *netSmooth* does, *netSmooth* outperforms them in amplifying the biological/technical variability ratio. *netSmooth* provides clusters that are more homogeneous and have higher adjusted mutual information (AMI) with respect to cell types. Although, in some cases data processed by MAGIC produces more robust clusters, the clusters returned after MAGIC processing do not have higher AMI or cluster purity. Higher robustness achieved by MAGIC processing might be due to the fact that the algorithm reinforces local structures too much in the data and producing artificially similar expression profiles between cells.

Finally, *netSmooth* is a versatile algorithm that may be incorporated in any analysis pipeline for any experiment where the organism in question has a high quality PPI network available. Although not shown, the algorithm is applicable to any omics data set that can be constructed as a genes-by-samples matrix, such as proteomics, SNPs and copy number variation. In addition, most of the computational load of network smoothing can be done “offline”. As such it scales well with the number of cells, which is likely to increase in future scRNA-seq experiments. We have made available an R package to that end, which is available on GitHub: https://github.com/BIMSBbioinfo/netSmooth.

## Methods and data

### The data sets

The hematopoiesis dataset [15] was obtained from the Gene Expression Omnibus [22]. The embryonic [18] and glioblastoma [19] datasets were obtained from *conquer* [23], a repository of uniformly processed scRNA-seq datasets.

### The random walks with restarts process

The *netSmooth* algorithm takes a graph *G* = {*V, E*} where *V* = {*gene_i_*} is the set of genes, and *E* = {(*i* → *j*)} is the set of edges between genes. The edge weights are degree-normalized, so that each gene’s outgoing edges’ weights sum to 1. We then define a process of random walk with restarts as in [13], on the PPI graph, where a conceptual random walker starts on a node in the graph (a gene/protein) and at each step walks to an adjacent node with the probability determined by the *α* times the edge weight. Further, at each step, there is a probability of (1 – *α*) that the walker restarts to its original node. Mathematically, given a graph defined by an adjacency matrix *A*[*_MxM_*], where *A_ij_* is the edge weight between gene *i* and gene *j* (and 0 for unconnected genes), and a vector *f*[*_Mx_*1], where *f_i_^t^* is the probability that the walker is at node *i* at step *t*, the process is defined by

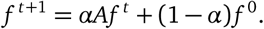

This process is convergent, and the stationary distribution is given by

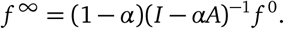

Hence, the random walk with restarts process is a diffusion process defined on the PPI graph, or through the diffusion kernel (smoothing kernel)

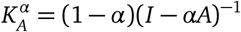

where (1 – *α*) is the restart probability, and *A* is the (column normalized) adjacency matrix of the PPI graph. Consequently, we define the *network-smoothed* expression profile

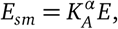

where *E*[_*MxN*_] is the normalized count values of the *M* genes in the *N* cells.

### The clustering procedure

Clustering analysis features prominently in scRNA-seq analyses; whether recapitulating known results or discovering new cell types, clustering cells by their gene expression profiles is commonly used to identify distinct populations. While some approaches directly take into account the zero-inflation of scRNA-seq data [5], other studies use traditional methods [18]. There is no standard method for clustering single cell RNAseq data, as different studies produce data with different topologies, which respond differently to the various clustering algorithms.

In order to avoid optimizing different clustering routines for the different datasets we benchmark on, we have implemented a robust clustering routine based on *clusterExperiment* [16], a framework for robust clustering based on consensus clustering of clustering assignments obtained from different clustering algorithms, different parameters for these algorithms, and different views of the data. The different views are different reduced dimensionality projections of the data based on different techniques. Thus, no single clustering result will dominate the data, and only cluster structures which are robust to different analyses will prevail. The procedure we implemented using the framework is as follows:

1. Perform different dimensionality reduction techniques on the data
  - PCA on the 500 most variable genes
    with 5 components
    with 15 components
    with 50 components
  - Alternatively to PCA, t-SNE on the 500 most variable genes
    with 2 dimensions
    with 3 dimensions
  - Select the most variable genes
    100 most variable genes
    500 most variable genes
    1000 most variable genes
2. On each reduced dimension view of the data, perform PAM clustering with K ranging from 5 to 10
3. Calculate the co-clustering index for each pair of samples (the proportion of times the samples are clustered together, in the different clustering results based on the different reduced dimensions and clustering parameters above)
4. Find a consensus clustering from the co-clustering matrix. This is done by constructing a dendrogram using average linkage, and traversing down the tree until a block with a self-similarity of at least 0.6, and a minimum size of 20 samples emerges. (instead of using cutree).
5. Perform hierarchical clustering of the cluster medioids, with similarities based on expression of the 500 most variable genes
6. Perform a DE analysis between clusters that are adjacent in the hierarchy from (5), and merge them if the proportion of genes that are found to be significantly differentially expressed between them (adjP <.05) is less than than 0.1.

Using only the 500 most variable genes insures the biological variation will dominate the technical variation, and enhances the reproducibility of t-SNE [24].

Importantly, samples that at step (4) don’t have a high enough affinity to any emerging cluster, will not be assigned to any cluster. The clustering is performed using the clusterExperiment::clusterSingle and clusterExperiment::clusterMany functions, the consensus clustering is obtained using the clusterExperiment::combineMany function, and the cluster merging (steps 5 and 6) using the clusterExperiment::makeDendrogram and clusterExperiment::mergeClusters functions. For more details, see [16].

### Choice of dimensionality reduction technique in the clustering procedure

In step (1) above, we cluster cells in a lower dimension embedding using either PCA [25] or t-SNE [26], in a dataset-dependent manner. Different single cell datasets respond better to different dimensionality reduction techniques which are better able to tease out the biological cluster structure of the data. In order to pick the right technique algorithmically, we compute the entropy in a 2D embedding. We obtained 2D embeddings from the 500 most variable genes using either PCA or t-SNE, binned them in a 20x20 grid, and computed the entropy using the discretize and entropy functions in the *entropy* R package [27]. The entropy in the 2D embedding is a measure for the information captured by it. For the clustering procedure, we pick the embedding with the highest information content. For the hematopoiesis and glioblastoma datasets, this is t-SNE, while for the embryonic development dataset it is PCA (Table 1). This method may be used to pick any dimensionality reduction technique other than the ones mentioned here, which might be more suitable for other analyses.

**Table 1.**
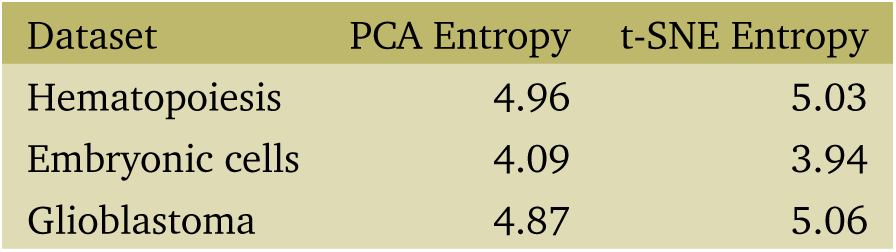
Entropy in 2D lower dimension embeddings

| Dataset | PCA Entropy | t-SNE Entropy |
| --- | --- | --- |
| Hematopoiesis | 4.96 | 5.03 |
| Embryonic cells | 4.09 | 3.94 |
| Glioblastoma | 4.87 | 5.06 |

### Cluster purity and adjusted mutual information

The cluster purity metric displayed above refers to the proportion of the samples in a cluster which are of the dominant cell type in that cluster. The purity for cluster *i* is given by

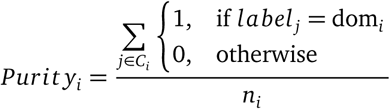

where *C_i_* = {*j*|cell_*j*_ ∈ cluster_*i*_}, *label_j_* is the cell type of *cell_j_*, *n_i_* = |*C_i_*| is the number of cells in cluster *i*, and

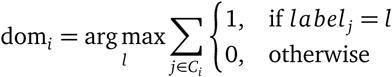

is the dominant cell type in cluster *C_i_*.

In addition to the cluster purity metric, we computed the Adjusted Mutual Information (AMI) [28], an information theoretic measure of clustering accuracy which accounts for true positives (two cells of the same type in the same cluster) being caused by chance. The AMI between a clustering *C* and the true labels *L* is given by

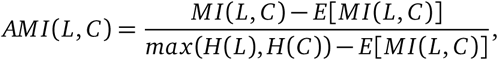

where *MI* (*a, b*) is the mutual information between labellings *a* and *b*, *H*(*a*) is entropy of clustering *a*, and *E*[·] denotes the expectation.

We do not compare the clusterings using the Rand index, as that measure penalizes for so-called *false negatives* (two cells of the same cell type but in different clusters), which is undesirable as cells from the same cell type might be rightly split into several clusters when a novel cell type is identified.

### Construction of the smoothing kernel

The PPI graph from which the diffusion kernel was derived was constructed using data from string-db [10]. For each pair of proteins, string-db provides a *combined interaction score*, which is a score indicating how confident we can be in the interaction between the proteins, given the different kinds of evidence string-db collates. We subset the links to only those above the 90th percentile of combined interaction scores, only keeping the 10% most confident interactions. For mouse that is 1,020,816 interactions among 17013 genes. For human, 852,722 interactions among 17467 genes.

### MAGIC and scImpute parameters

For all the results presented in this paper, scImpute was run using the default parameters (drop_thre = 0.5). For MAGIC, we used values for the diffusion time parameter (*T* = {1,2,4,8,16}). Unlike *netSmooth,* for MAGIC the proportion of samples in robust clusters and the cluster purities were anti-correlated; thus we picked the one that gave the best cluster purities as the best MAGIC parameter. The chosen T values are given in Table 2.

**Table 2.**
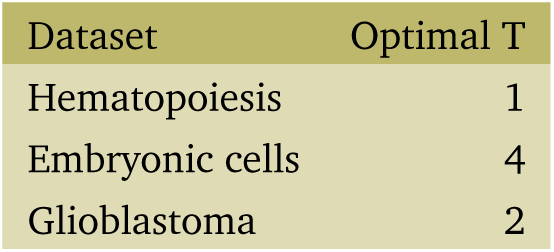
Opitimal diffusion time values for MAGIC.

| Dataset | Optimal T |
| --- | --- |
| Hematopoiesis | 1 |
| Embryonic cells | 4 |
| Glioblastoma | 2 |

### the *netSmooth* R package

The analysis for this paper was done using the companion *netSmooth* R-package, which is available online: https://github.com/BIMSBbioinfo/netSmooth.

## Footnotes

1 Most interactions in string-db do not specify the direction, or nature of the interaction

## Author contributions

AA conceptualized the project, AA and JR conceived of the algorithm together. All the analysis and software development was done by JR, who also wrote the initial draft of manuscript with input from AA. AA supervised the writing, software development and analysis. JR wrote R package with input and code review and contributions from AA.

## Acknowledgements

We would like to thank Vedran Franke, Bora Uyar and Brendan Osberg for valuable comments and input for the development of this manuscript.

## Supplementary material

**Figure S1.**
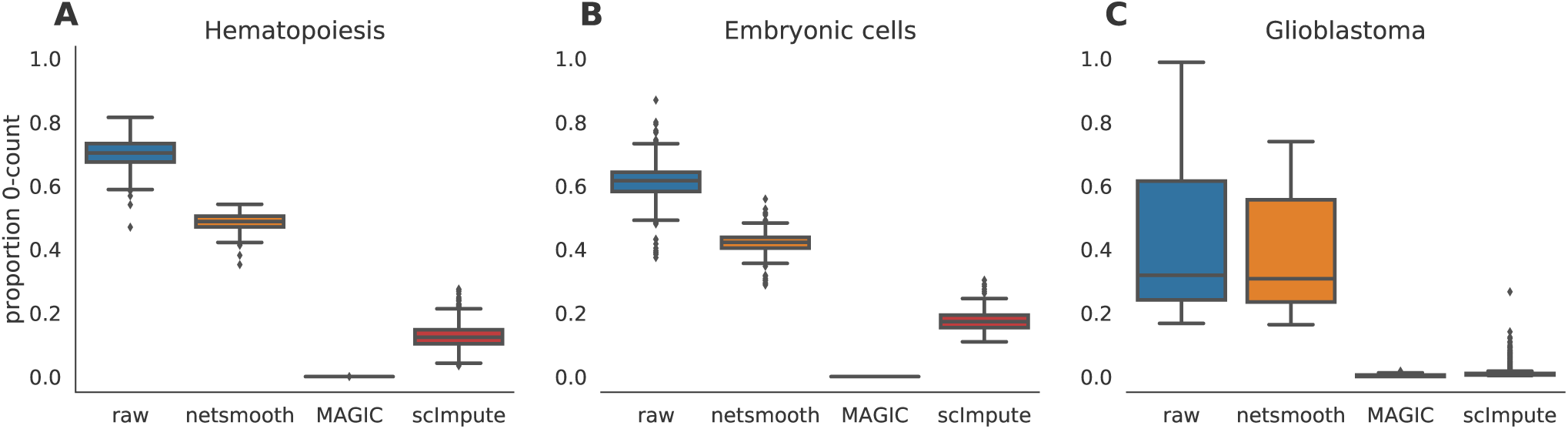
The proportion of genes with 0 counts is a proxy for technical dropouts. A) no preprocessing, B) after application of *netSmooth*, C), using scImpute, and D) after application of MAGIC.

**Figure S2.**
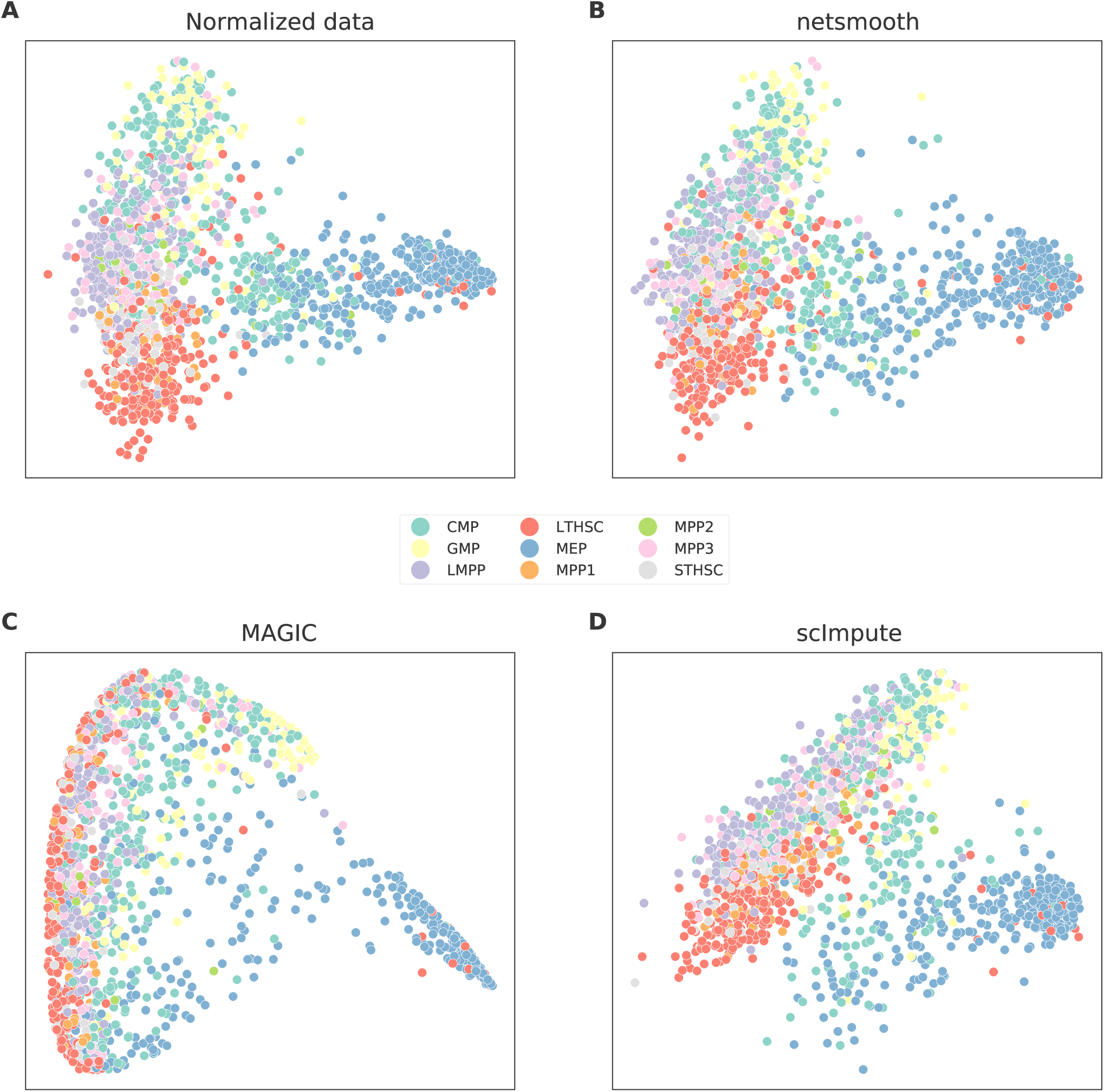
PCA plots of the HSPC dataset A) no preprocessing, B) after application of *netSmooth*, C), using scImpute, and D) after application of MAGIC.

**Figure S3.**
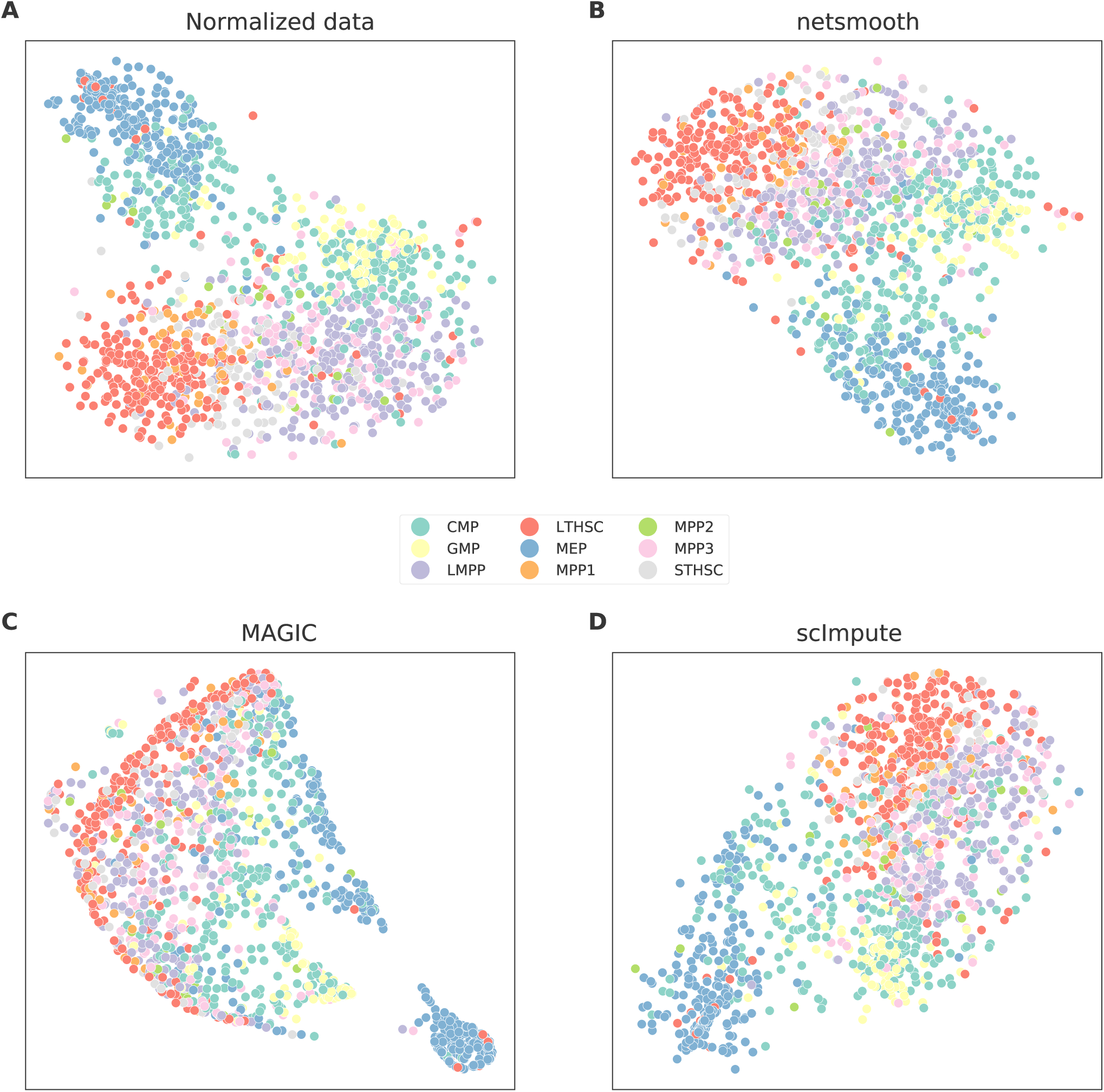
t-SNE plots of the HSPC dataset A) no preprocessing, B) after application of *netSmooth*, C), using scImpute, and D) after application of MAGIC.

**Figure S4.**
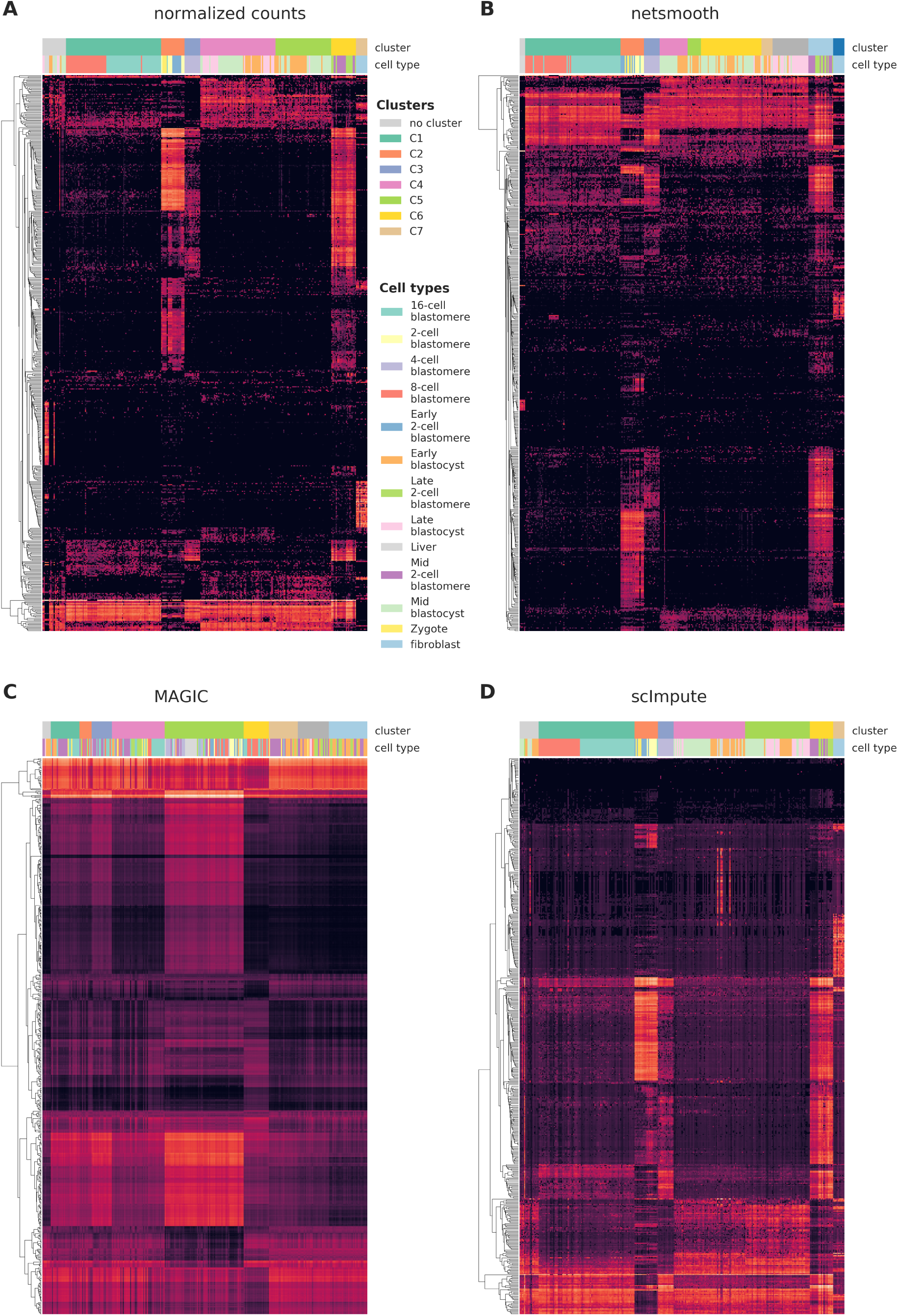
single cells from the embryonic development dataset were clustered using the robust clustering procedure, and the 500 most differentially expressed genes (by edgeR-QLF test adjusted P value) in any of the discovered clusters are shown in a heatmap, as well as cluster assignments and cell types. A) raw (no imputation), B) after application of *netSmooth*, C) missing values imputed using scImpute D) after application of MAGIC.

**Figure S5.**
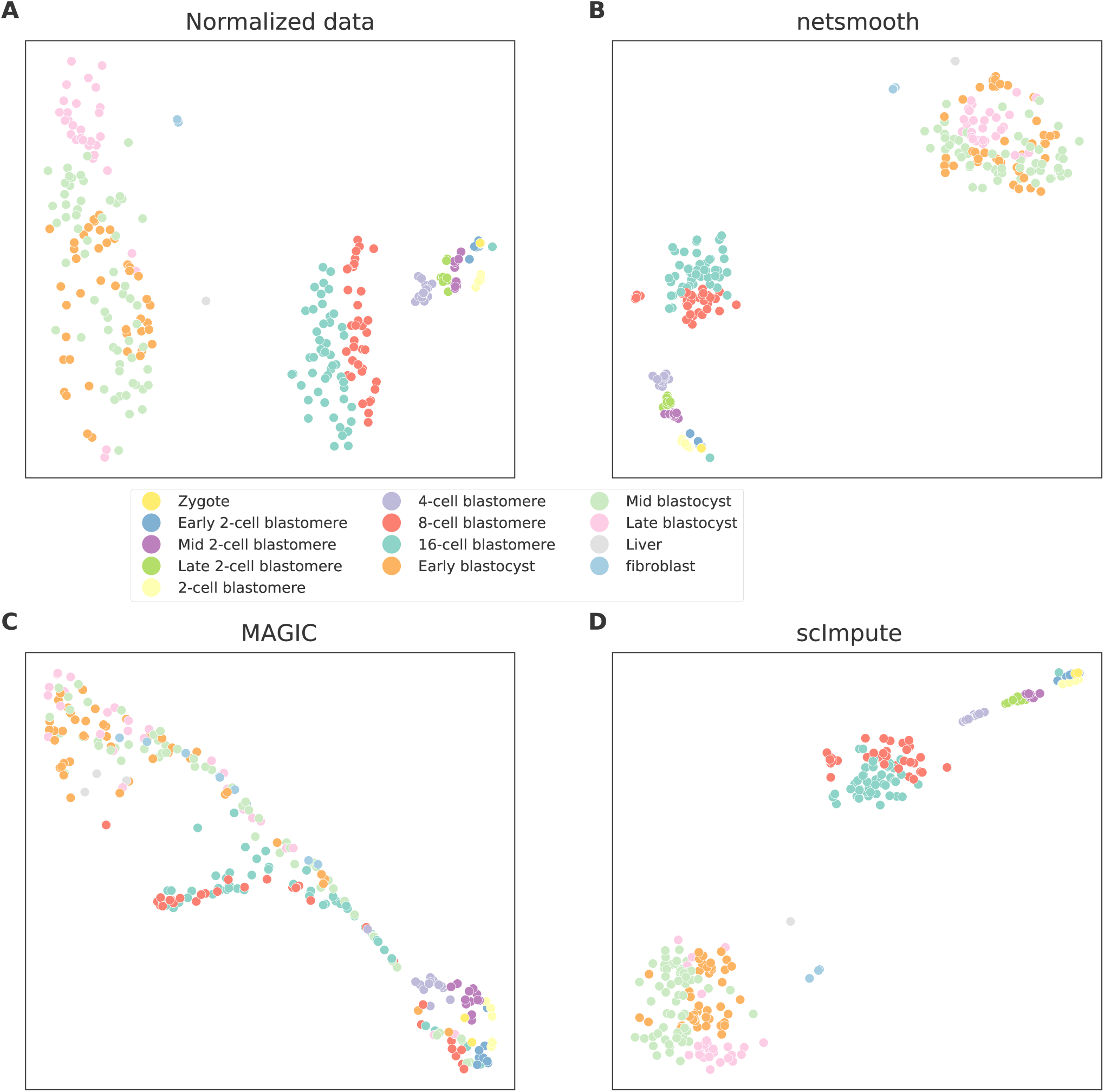
t-SNE plots of the embvryonic development dataset A) no preprocessing, B) after application of *netSmooth*, C), using scImpute, and D) after application of MAGIC.

**Figure S6.**
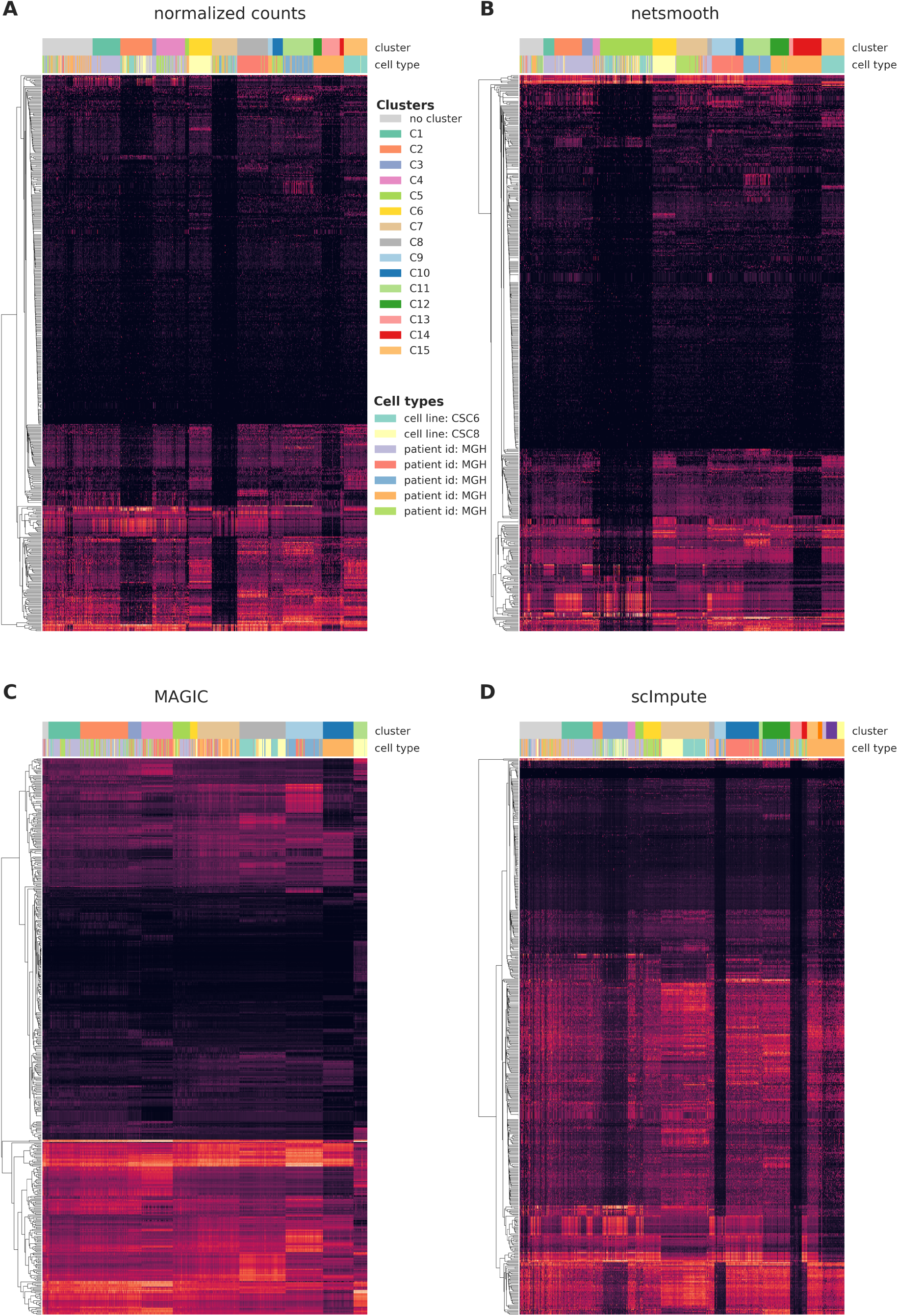
single cells from the glioblastoma dataset were clustered using the robust clustering procedure, and the 500 most differentially expressed genes (by edgeR-QLF test adjusted P value) in any of the discovered clusters are shown in a heatmap, as well as cluster assignments and cell types. A) raw (no imputation), B) after application of *netSmooth*, C) missing values imputed using scImpute D) after application of MAGIC.

**Figure S7.**
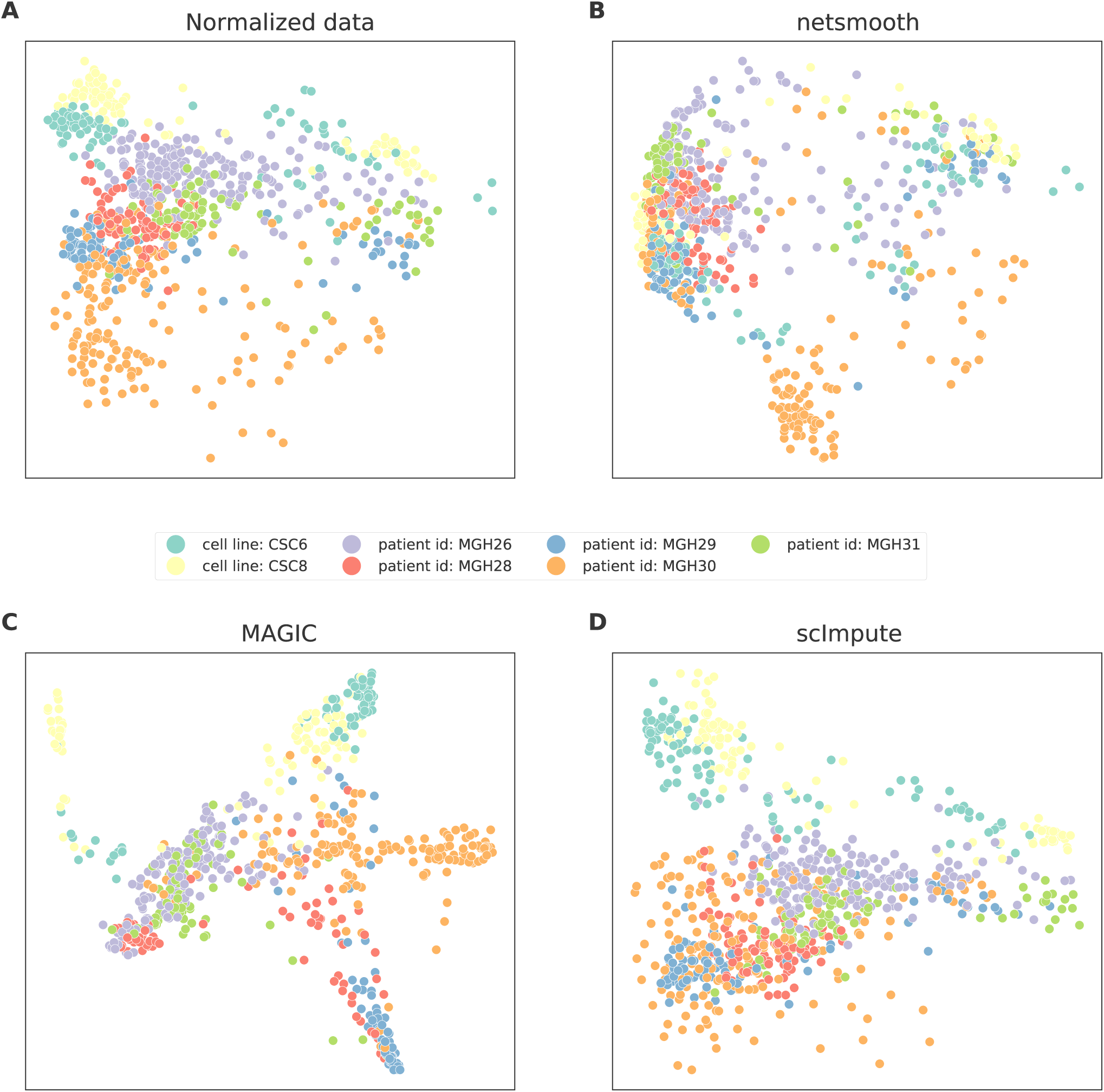
PCA plots of the glioblastoma dataset A) no preprocessing, B) after application of *netSmooth*, C), using scImpute, and D) after application of MAGIC.

**Figure S8.**
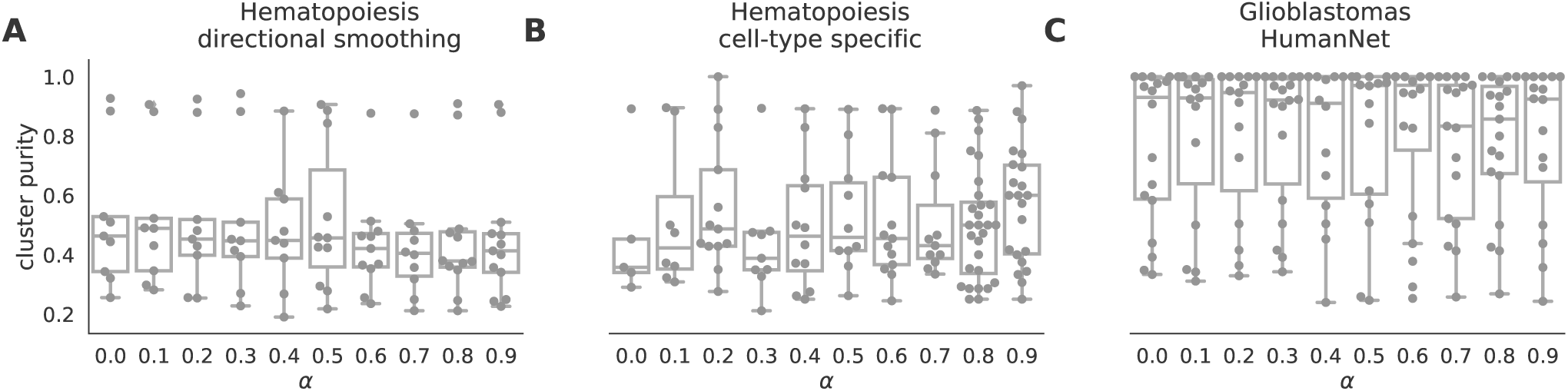
Cluter purity by smoothing parameter. A) for the hematopoiesis dataset with a directional (signed) graph, where inhibitory interactions have a negative edge weight. B) For the hematopoiesis dataset using a gene network with only genes that have a cell-type specific expression in any cell type. C) In the glioblastoma dataset using a gene network from HumanNet.

